# Controlling noisy expression through auto regulation of burst frequency and protein stability

**DOI:** 10.1101/511774

**Authors:** Pavol Bokes, Abhyudai Singh

## Abstract

Protein levels can be controlled by regulating protein synthesis or half life. The aim of this paper is to investigate how introducing feedback in burst frequency or protein decay rate affects the stochastic distribution of protein level. Using a tractable hybrid mathematical framework, we show that the two feedback pathways lead to the same mean and noise predictions in the small-noise regime. Away from the small-noise regime, feedback in decay rate outperforms feedback in burst frequency in terms of noise control. The difference is particularly conspicuous in the strong-feedback regime. We also formulate a fine-grained discrete model which reduces to the hybrid model in the large system-size limit. We show how to approximate the discrete protein copy-number distribution and its Fano factor using hybrid theory. We also demonstrate that the hybrid model reduces to an ordinary differential equation in the limit of small noise. Our study thus contains a comparative evaluation of feedback in burst frequency and decay rate, and provides additional results on model reduction and approximation.

## 1 Introduction

Synthesis of protein molecules in bursts of multiple copies has been identified as a major factor in gene expression noise [12]. The number of bursts per protein lifespan determines the abundance of a bursty protein [9]. This ratio can be controlled by the numerator, the burst frequency, or the denominator, the protein decay rate. Feedback in burst frequency has been widely documented [2], and examples of feedback in decay rate are available too [15, 26]. Linear noise approximation based analysis suggest that the two feedback pathways are equivalent in terms of controlling gene-expression noise [25].

In this paper we compare the two feedback pathways using a hybrid model for bursty gene expression with negative feedback in burst frequency or decay rate. Hybrid models mix continuous deterministic with discrete stochastic dynamics [10, 11, 18, 22]. The chosen modelling framework is hybrid in that it combines stochastic dynamics of bursty production occurring at discrete time-points with deterministic dynamics of protein decay [4].

Intuition suggests that, by repressing production, negative feedback lowers the protein mean, and, by improving regression to the mean, it also lowers the protein noise [13]. Counter-intuitively, multiple studies report that the response of noise to increasing feedback strength is U-shaped [19, 27]. The eventual increase in the noise can be attributed e.g. to low copy number effects [24], loss of time averaging [17], or the failure to control large bursts [5]. In this paper we examine how the choice of feedback pathway (burst frequency or decay rate) affects the shape of the noise response to strengthening feedback.

The outline of the paper is as follows. Section 2 introduces the chosen hybrid modelling framework on a protein which is expressed constitutively without feedback. Section 3 extends the hybrid model by negative autoregulation, and Section 4 derives the steady-state distribution for the extended model. Section 5 defines a specific noise metric that is used here to evaluate feedback performance. Sections 6 and 7 elaborate on feedback in burst size and decay rate, respectively, the two feedback pathways whose performance we are set to compare. Section 8 introduces a full discrete model for bursty protein expression. Section 9 contains the bulk of the results of this paper that are based on the theoretical backbone of the previous sections. The results compare the performance of the two feedback types, and draw connections between the full discrete, the hybrid and the deterministic modelling formalisms. Section 10 concludes the paper with a short summary.

## 2 Constitutive model

By the constitutive model we understand a hybrid stochastic bursting gene-expression model without a feedback mechanism. The protein level dynamics is given by the balance of deterministic protein decay and stochastic protein synthesis in bursts. Between bursts, the protein concentration satisfies a linear ordinary differential equation d*x*/d*t* = −*γx*, where *γ* is the decay rate constant, implying that the temporal profile of protein concentration is piecewise exponential (see Fig 1, left). Bursts occur randomly in time with frequency *α* per unit time. It follows that the waiting time from one burst until the next one is drawn from the exponential distribution with mean waiting time 1/*α*. The size of a burst is also random and is drawn from the exponential distribution with mean burst size *β* [9].

**Fig. 1.**
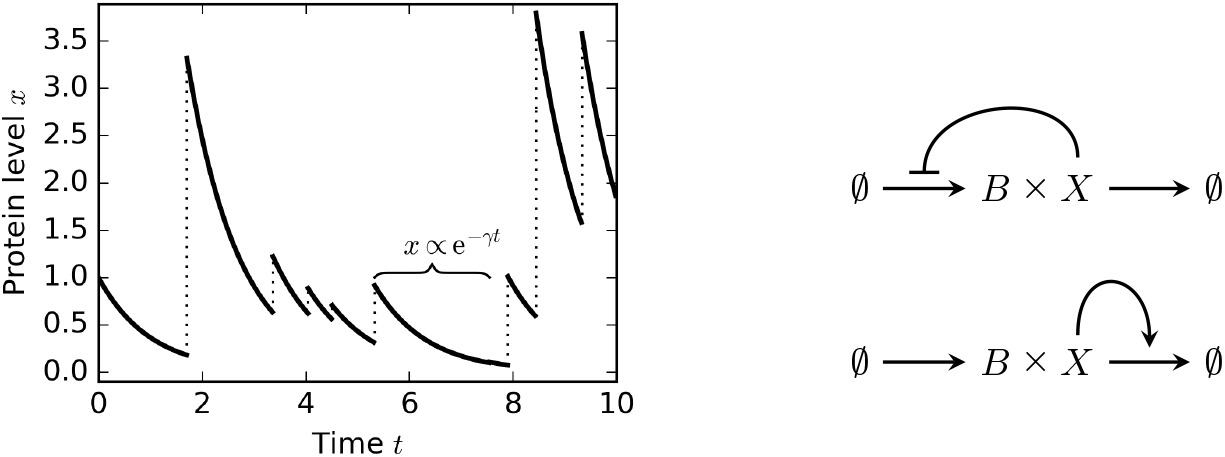
*Left:* Sample protein path corresponding to the constitutive model (1). The concentration of protein decays with rate constant *γ* between bursts (solid lines). Bursts occur randomly in time with burst frequency *α* and lead to positive discontinuous jumps of mean size *β* in protein level (dotted vertical lines). In this example, *α* = *β* = *γ* = 1. *Right:* The two feedback types considered in this paper are feedback in burst frequency and feedback in decay rate. Protein X is produced in bursts of size B and degrades one molecule at a time. The protein controls its level either by reducing the frequency of burst occurrence or by enhancing its own decay.

The master equation for the hybrid process as described above takes the form of a partial integro-differential equation [23]

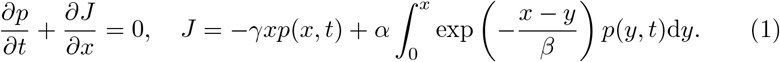

The solution *p*(*x*, *t*) gives the probability density function of protein concentration *x* at time *t*.

The first equation in (1) states the principle of probability conservation in differential form. It says that the probability changes in time due to differentials in the probability flux J. The probability flux is described in the second equation of (1). The flux consists of a negative local flux and a positive non-local flux. In general, a flux is local if it depends on the value of the solution at the point x of flux evaluation, whereas a non-local flux depends on the values of the solution away from the evaluation point; the sign of a flux corresponds to the direction of probability mass transfer. In our particular model (1), the negative local flux represents the downward transfer of probability mass due to deterministic decay of protein, and the positive non-local flux represents the upward transfer of probability mass due to bursts of protein production. Note in particular that the integral kernel in the nonlocal flux expresses the probability that a burst occurs which takes the protein concentration from a value *y* below *x* into any value above *x*.

Previous studies established that the gamma distribution [14]

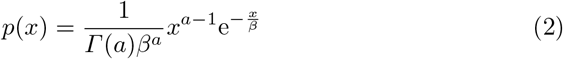

is a steady-state solution to the master equation (1). The parameter *a* in (2) is defined by

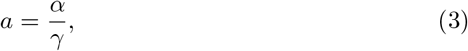

and gives the average number of protein bursts per protein lifetime. The mean and variance of (2) are

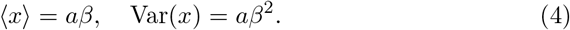

It follows immediately from (4) that the squared coefficient of variation, defined as the ratio of the variance to the square of mean, is equal to *a*^−1^. Therefore, if a large number of bursts occur on average per protein lifetime, the noise in protein concentration is low.

It is convenient to measure the protein concentration in units of its mean and time in units of the protein lifetime. This is achieved via nondimensionalisation

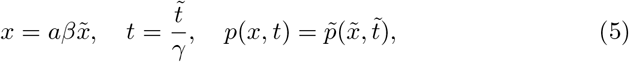

where 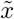 and 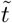 represent the dimensionless concentration and time variables. We insert (5) into (1) and, for simplicity, drop the tildes in the dimensionless variables symbols, obtaining

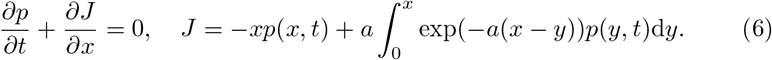

Comparing the dimensional problem (1) to the dimensionless problem (6), we see that the latter can be formally obtained from the former by setting *α* = *a*, *β* = 1/*a*, *γ* = 1. However, the assignment *β* = 1/*a* should not be interpreted as implying that burst sizes are physically small if the burst frequency is large. Rather, it means that burst sizes are small in comparison to the steady-state mean.

In the regime *a* ≫ 1 of very frequent (and very short) bursts, the probability flux *J* in (6) can be approximated using the Laplace method [16] by

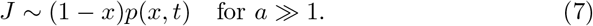

The reduced flux (7) corresponds to deterministic dynamics governed by the ordinary differential equation

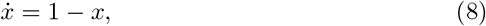

whose solutions have the explicit form of

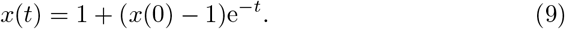

Figure 2 compares sample paths of the hybrid model (6) to the deterministic solution (9) for two different initial conditions, *x*(0) = 0 (Fig 2, left) and *x*(0) = 2 (Fig 2, right). As expected from the use of the Laplace method, stochastic sample paths are close to the deterministic solution for large burst frequencies *a*.

**Fig. 2.**
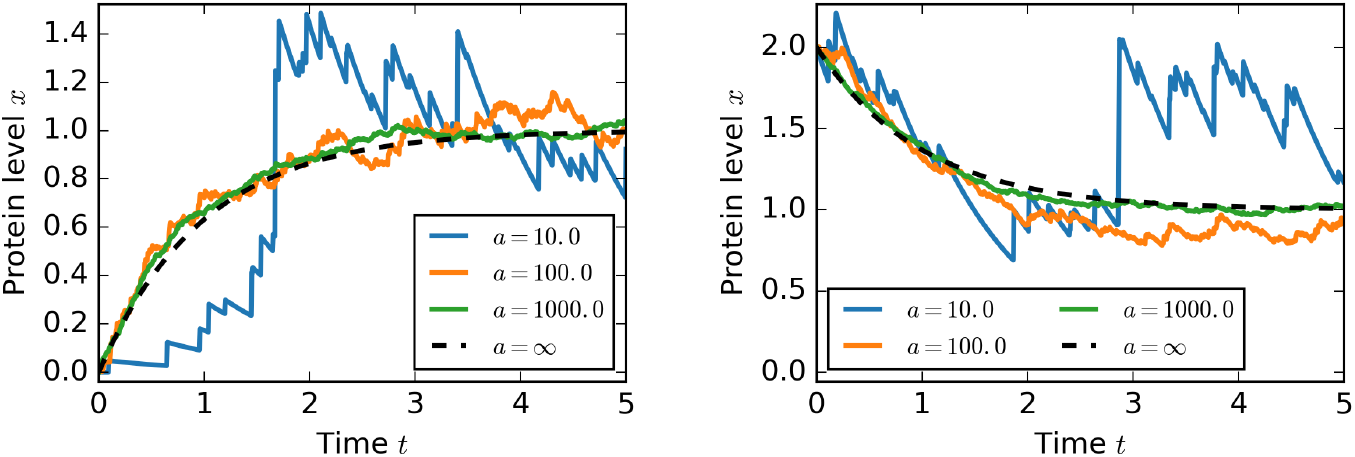
As the average number *a* of bursts per protein lifetime increases, the trajectories of the stochastic bursting model (6) become well approximated by the solution (9) to the ordinary differential equation model (8). Without loss of generality, the burst size is scaled to 1/*a* in order that the steady-state protein mean is equal to 1 regardless of the choice of *a*.

## 3 Feedback model

Here we extend the hybrid stochastic model (6) with feedback in burst frequency and decay rate (Fig 1, right). In the feedback model, the probability of a burst to occur in a time interval of length dt is equal to *ah*(*x*)*dt* + *o*(*dt*), where *x* gives the current protein concentration and *h*(*x*) is a response function as specified below. Bursts sizes are exponentially distributed with mean size 1/*a* like in the (dimensionless) constitutive model. Between bursts, the protein concentration satisfies 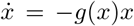, in which *g*(*x*) is another response function. We assume that the response functions satisfy

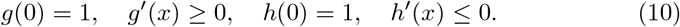

If there is a shortage of protein (*x* is close to zero), bursts occur with frequency *a* and decay with rate constant 1 as in the constitutive model. However, as the protein concentration increases, bursts become less frequent and/or the propensity of protein molecules for decay increases.

The master equation for the feedback model reads

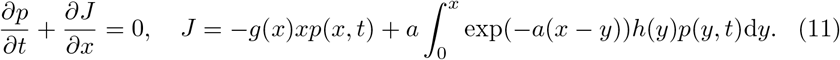

Applying the Laplace method on the non-local flux yields

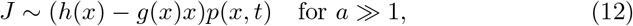

which corresponds to the ordinary differential equation

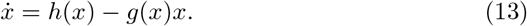

Under assumptions (10), equation (13) has a single globally stable steady state which is smaller than the steady state 1 of the constitutive deterministic model (8).

## 4 Steady state distribution and moments

At steady state, the probability flux *J* in the master equation (11) vanishes, leading to the Volterra integral equation

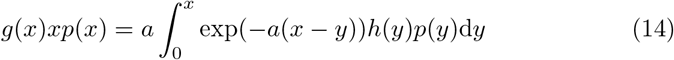

for the stationary protein distribution *p*(*x*). In order to solve (14) in the unknown *p*(*x*), we multiply both sides by e^*ax*^,

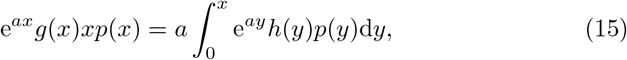

and differentiate with respect to *x* to obtain a linear first-order ordinary differential equation

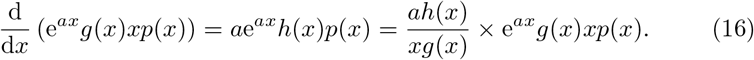

Solving (16) in e^*ax*^ *g*(*x*)*xp*(*x*) implies that up to a normalisation constant we have

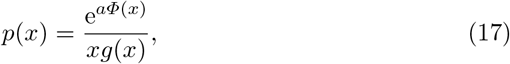

in which the potential *Φ*(*x*) is defined through the indefinite integral

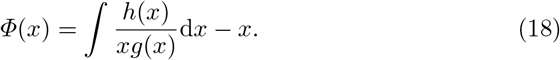

The *n*-th moment of the steady-state protein distribution is given by

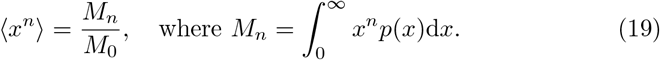

In general, the moments can be evaluated by numerical integration of (19). In special parametric regimes, asymptotic approximations to the integrals (19) can be developed (Appendices A and B). In the next section, we use the moments (19) to define a specific characteristic of protein noise.

## 5 Relative noise

In this section we provide a definition of relative noise in protein concentration. The purpose of this quantity is to compare the steady-state variance in a feedback model to the steady-state variance in a referential constitutive model. The latter is chosen so as to have the same steady-state mean as the feedback model. By doing such a comparison, we compensate for the increase in noise in the feedback model that results from the decrease of the time-averaged number of bursts per protein lifetime. What remains is the change in noise that results from improved mean reversion in a feedback model. Indeed, we shall see that the relative noise is always less than 1 in our examples of negative autoregulatory pathways. For this section only, we refer to the concentration of a self-regulating protein as *x*_reg_ and to the concentration of the referential constitutive protein as *x*_const_.

In the absence of regulation, the normalised burst frequency is equal to *a* and the burst size is equal to 1/*a*. These values lead to the mean value of 1. In order to satisfy the constraint 〈*x*_const_〉 = 〈*x*_reg_〉, we decrease the normalised burst frequency in the constitutive model to *a*〈*x*_reg_〉 whilst keeping the burst size equal to 1/*a*. Since the protein variance is equal to the product of burst frequency and the square of burst size, cf. (4), we find that

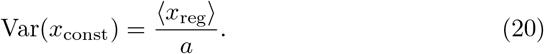

The relative noise compares the variances in the regulated model and the referential constitutive model,

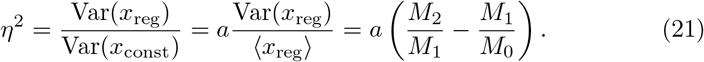

The definition (21) of the relative noise superficially resembles the Fano factor [6]. However, the two should not be confused. The value one of Fano factor means Poissonian noise. On the other hand, *η*^2^ = 1 means that the regulated protein has the same variance as the referential unregulated protein. Nevertheless, that can still correspond to a very large Fano factor: how large the actual Fano factor is depends on how many molecule copies are encompassed in an average burst. In Section 8, we consider a discrete modelling approach and systematically establish the relationship between the Fano factor of a full discrete model and the relative noise of the hybrid model.

## 6 Feedback in burst frequency

Sections 3–4 provided general results for feedback in burst frequency and decay rate acting concurrently. Here we provide additional details for the situation if feedback is in burst frequency only. We thereby focus on a specific type of response function, the decreasing Hill function. This leads to choices

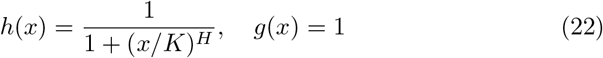

in the general model (11). The parameter *K* gives the critical concentration of protein that is required to halve the burst frequency. The parameter *H* is the cooperativity coefficient. Large values of *H* imply that the burst freuquency decreases rapidly from its maximal value to zero as the protein concentration exceeds the critical threshold *K*. The critical threshold *K* is a reciprocal measure of feedback strength: small values of *K* mean that low amounts of protein suffice to turn off the production. For this reason we refer from now on to the reciprocal *K*^−1^ of the critical threshold as feedback strength. It is easy to verify that the choices in (22) satisfy the assumptions (10) imposed on the feedback model.

With choices (22), the limiting ordinary differential equation (13) takes the form of

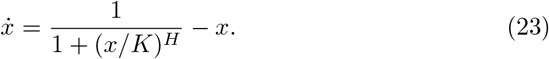

Solutions to (23) converge to a unique, globally stable, steady state, which satisfies the fixed-point equation

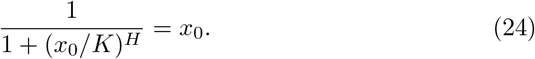

Elementary analysis shows that *x*_0_ is an increasing function of *K*, i.e. that the deterministic steady state *x*_0_ decreases with increasing feedback strength *K*^−1^. With choices (22), the potential (18) is an elementary function

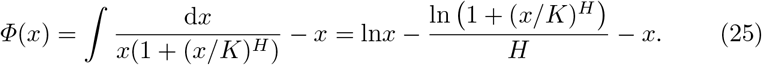

Inserting (25) into (17) we find an explicit formula

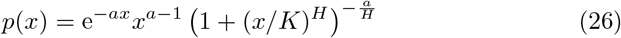

for the steady-state protein pdf which holds up to a normalisation constant. The asymptotic behaviour of the mean 〈*x*〉 (19) and the relative noise *η*^2^ (21) for the protein pdf (26) in the small-noise regime (*a* ≫ 1) and the strong feedback regime (*K* ≪ 1) is provided in Appendices A and B.

## 7 Feedback in decay rate

Here we explore in detail the situation if feedback is in decay rate only. Specifically, we use the choices

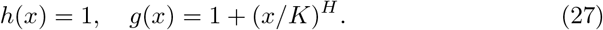

The polynomial response function *g*(*x*) consist of a basal term 1 and a monomial term which is proportional to *x^H^*. Biologically, this means that in addition to spontaneous decay, there is an additional decay pathway, which is cooperatively activated by the protein itself. The critical concentration *K* gives the amount of protein that is necessary to double the rate of decay per protein molecule. Small values of *K* mean that few proteins suffice to turn on the decay, suggesting that the reciptocal *K*^−1^ can again be used as a measure of feedback strength.

With choices (27), the limiting ordinary differential equation (13) reads

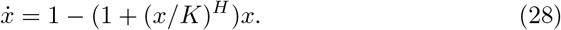

Equation (28) describes a different time-dependent dynamics than the limiting equation (23) for feedback in burst frequency. Nevertheless, solutions to (28) converge to the same steady-solution *x*_0_ satisfying (24). Furthermore, the probability-distribution potential (18) for the choices (27) is the same as (25) obtained for feedback in burst frequency.

The steady-state protein pdf (17) simplifies to

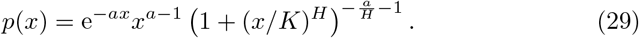

We note that the pdf (29) differs from (26) only in the exponent of the third factor. The asymptotic behaviour of the mean 〈*x*〉 (19) and the relative noise *η*^2^ (21) for the protein pdf (29) in the small-noise regime (*a* ≫ 1) and the strong feedback regime (*K* ≪ 1) is provided in Appendices A and B.

## 8 Discrete approach

The full discrete model for feedback in burst frequency and decay rate is based on chemical reactions

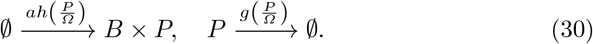

The first reaction in (30) is the production of protein *P* in bursts of size *B*. The second reaction in (30) is the degradation of protein *P*. The response functions depend on the ratio *P/Ω* of the protein copy number to a system-size parameter *Ω*. Large values of *Ω* mean that feedback is sensitive to large changes in protein molecules. As with the hybrid model, we treat the discrete model (30) separately for the choices (22) (feedback in burst frequency) and the choices (27) (feedback in decay rate).

The burst size *B* is assumed to be drawn from the geometric distribution [21] with mean *Ω/a*

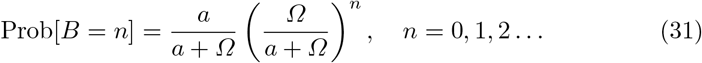

Due to previously developed theoretical arguments [3, 8, 20], the protein concentration *x* = *P*/*Ω* approximately satisfies in the large system-size limit *Ω* ≫ 1 the hybrid bursting model (11).

The Fano factor, which is defined as the variance to mean ratio, is a widely used measure of variability in discrete probability distributions and discrete stochastic models for gene expression [6]. The protein Fano factor satisfies

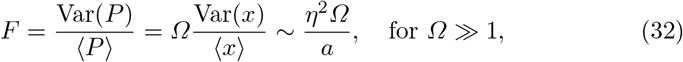

where *η*^2^ is the relative noise of the hybrid model as defined by (21). Hence, for large system sizes, the Fano factor is proportional to the mean burst size *Ω*/*a*, with the relative noise of the hybrid model giving the factor of proportionality.

## 9 Results

In this paper we explored a bursting model for stochastic gene expression with negative feedback. The model is hybrid in the sense that it combines a deterministic decay of protein with stochastic protein production in bursts. Two separate versions of the model were considered, depending on whether the feedback is in burst frequency or decay rate. The model (either version) is characterised by three parameters: the maximal burst frequency *a;* the critical concentration *K*; and the cooperativity coefficient *H*.

The maximal burst frequency *a* gives the average number of bursts per protein lifetime at full gene activation. Without loss of generality, the mean burst size is scaled to 1/*a*. This choice of scaling implies that the mean protein level, which is equal to the product of the burst frequency and the burst size, is bounded by one. In the regime *a* ≫ 1 of frequent bursts, the trajectories of the stochastic model fluctuate near the deterministic solution (Fig 2). The regime *a* ≫ 1 is therefore referred to as the small-noise regime.

The parameters *K* and *H* determine the character of the feedback response. The critical concentration *K* gives the amount of protein that is required to halve the frequency of bursts (in case of feedback in burst frequency) or double the propensity for decay (in case of feedback in decay rate). The reciprocal *K*^−1^ is used as a measure of feedback strength. The regime *K* ≪ 1 (i.e. *K*^−1^ ≫ 1) is referred to as the strong-feedback regime of the model. The cooperativity coefficient *H* determines how steeply the response changes as the protein concentration passes through the critical threshold *K*.

Figure 3 shows the steady-state values of protein mean and relative noise as functions of feedback strength. The relative noise is defined in Equation (21) as the ratio of the variance of the protein with feedback to the variance of a constitutively expressed protein with the same mean. Several values of the maximal burst frequency *a* are selected, including the limit value of *a* → ∞, the results for which are derived in Appendix A using a small-noise approximation. The cooperativity coefficient is set to *H* = 4.

**Fig. 3.**
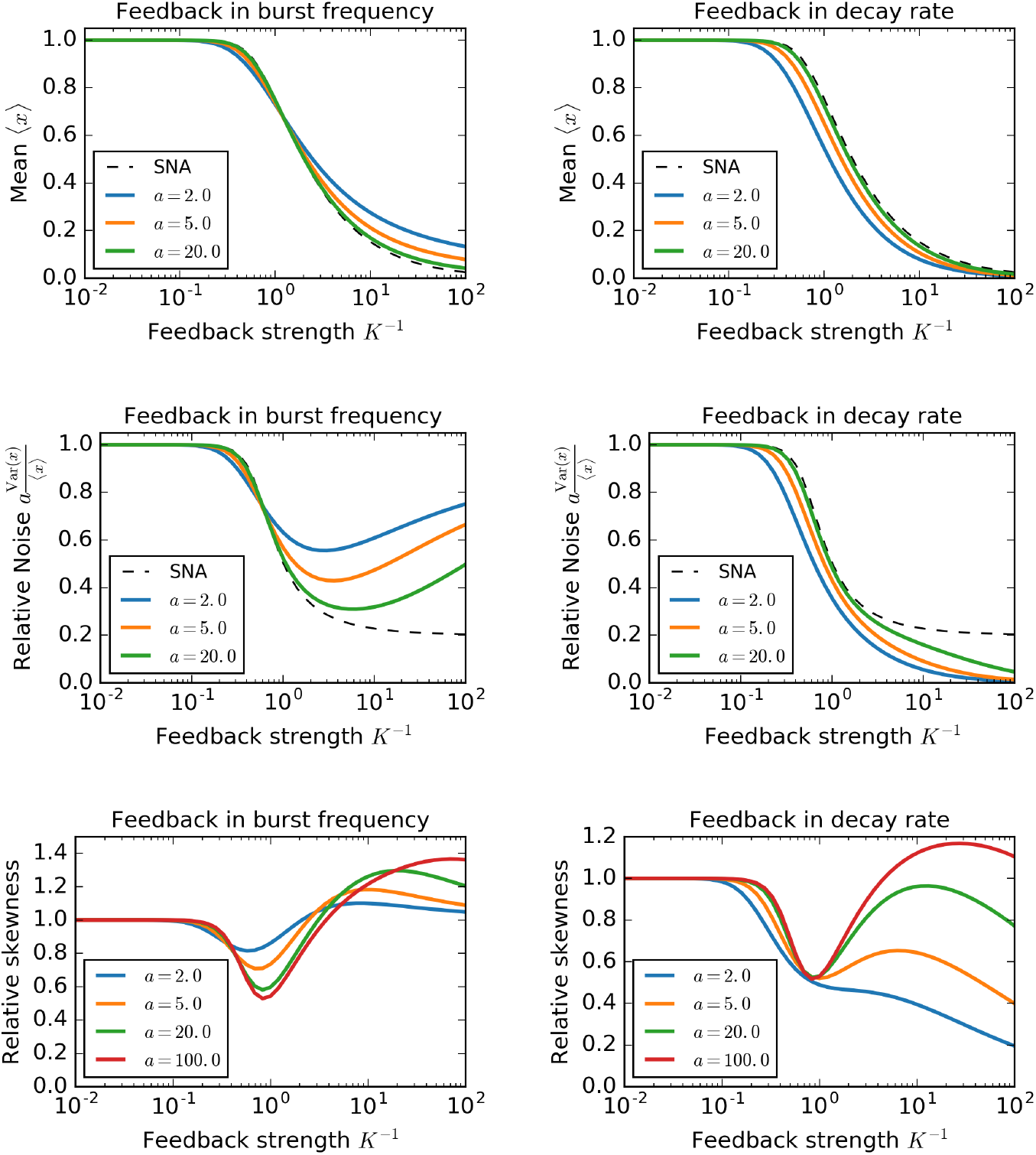
The mean, the relative noise and skewness of protein concentration subject to feedback in burst frequency or decay rate. The exact values (coloured lines) are based on numerical integration of (19), in which the probability density function *p*(*x*) is given by (26) (feedback in burst frequency) or (29) (feedback in decay rate). The small-noise approximation (SNA) of the protein mean is the fixed-point solution xo to Equation (24); the SNA of the relative noise is given by (A4). The feedback cooperativity coefficient is fixed to *H* = 4 throughout.

The small-noise approximation leads to the same mean and noise values for both feedback types. In particular, the small-noise approximation suggests that, regardless of the feedback type, a maximal (*H* + 1)-fold reduction of relative noise can be achieved in the limit *K* → 0 of strong feedback. However, the assumption of high burst frequency, on which the use of small-noise approximation is based, eventually breaks down as feedback strengthens. Indeed, finite values of the maximal burst frequency *a* paint a radically different picture from that obtained by the small-noise approximation. In case of feedback in burst frequency, the relative noise starts increasing for large feedback strengths, eventually returning to the value of one. Contrastingly, in case of feedback in decay rate, the relative noise decreases down to zero. Thus, despite the small-noise prediction that the two feedbacks are indistinguishable in terms of controlling protein mean and noise, we see that at high feedback strengths, feedback in decay rate can be much more effective than feedback in burst frequency.

The stark differences between the small-noise prediction and the exact results motivated us to develop in Appendix B an alternative asymptotic approximation in the regime *K* ≪ 1 of strong feedback. For feedback in decay rate the asymptotics (B6) reveal that the relative noise is: of the order of *K* if *H* > 2; of the (asymptotically larger) order of *K*^*H*−1^ if 1 < *H* < 2; or converges to the constant 1 − *H* if 0 < *H* < 1. For feedback in burst frequency the strong-feedback asymptotics (B7) confirm the numerical observation that the relative noise eventually returns to the value of one as *K* tends to zero.

The protein skewness is quantified by the third standardised moment of its steady-state distribution. In the nethermost panels of Fig 3 we report the response of a relative protein skewness to increasing feedback strength. By the relative skewness we understand the ratio of the skewness of the auto-regulated protein to that of a referential constitutive protein with the same mean. The protein skewness responds to increasing feedback strength in a complicated manner featuring first a trough and then a peak. Feedback in burst frequency is more conducive to skewness than feedback in decay rate. Details on the mathematical definition and calculation of the (relative) skewness are provided in the Appendix C.

In order to cross-validate the hybrid framework, we constructed a fine-grained discrete stochastic bursting model (30) with feedback in burst frequency or decay rate. In the discrete model, burst sizes are geometrically distributed; decay is stochastic and leads to the removal of one molecule at a time. In addition to the three parameters of the hybrid model, the discrete model features an additional system-size parameter *Ω*, which is equal to the copy number *P* of protein that are encompassed in a unit of protein concentration *x*. Provided that *Ω* is large, discrete protein distributions obtained by stochastic simulation of the discrete model (30) are well approximated by the explicit continuous protein distributions (26) or (29) obtained using hybrid theory (Fig 4).

**Fig. 4.**
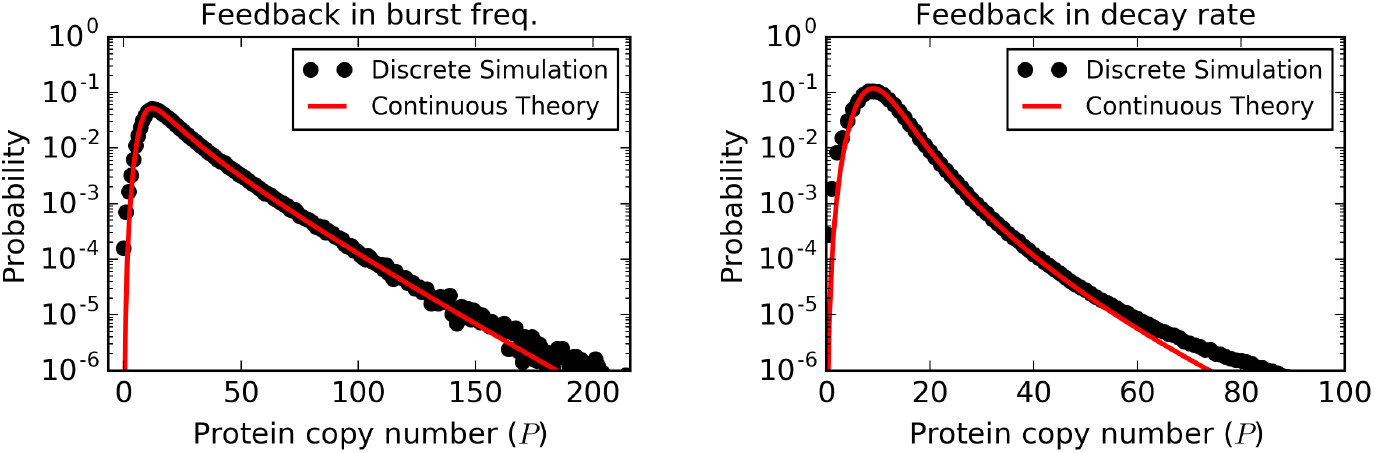
Steady-state protein copy number distribution by discrete simulations and hybrid (continuous) theory. The theoretical distribution is given by 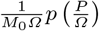, where *p* is the continuous probability density function (26) (feedback in burst frequency) or (29) (feedback in decay rate) of the hybrid model and *M*_0_ is the zero-th moment (19) (the reciprocal of the normalisation constant). Discrete simulation results are based on 10^6^ Gillespie iterations of the discrete model (30), in which the response functions *h*(*x*) and *g*(*x*) are given by (22) (feedback in burst frequency) or (27) (feedback in decay rate). The model parameters are: burst frequency *a* = 5; cooperativity coefficient *H* = 4; critical concentration *K* = 0.1; system size *Ω* = 100.

The variability of a discrete distribution is conveniently quantified using the Fano factor (the variance to mean ratio). Figure 5 shows the Fano factor for the steady-state protein copy number obtained by stochastic simulation of the full discrete model (30) and the hybrid theory approximation (32). For large values of *Ω* the two agree well. For small values of *Ω*, single-molecule effects become important in discrete simulations; by neglecting them, the hybrid theory tends to underestimate the Fano factor.

**Fig. 5.**
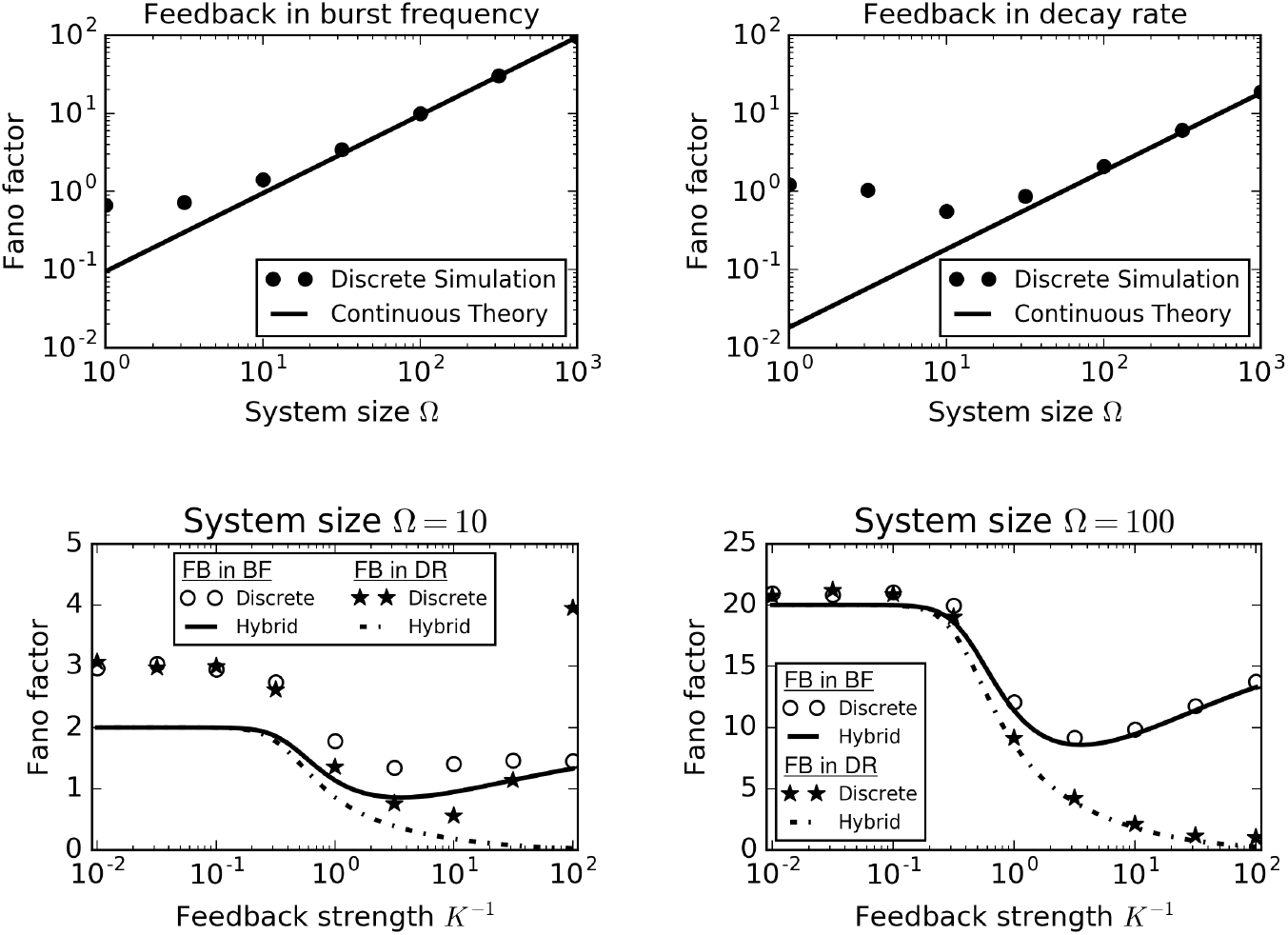
The Fano factor of steady-state protein copy number as function of system size *Ω* (upper panels) and feedback strength *K*^−1^ (lower panels). The discrete results are obtained by simulation of system (30). The hybrid (continuous) are based on (32), (21), and (19). The model parameters are set to *a* = 5, *H* = 4, *K* = 0.1 (upper panels) *Ω* = 10 or *Ω* = 100 (lower panels). The number of Gillespie iterations is set to 10^4^ × ⌊*Ω*⌋.

The hybrid approximation (32) implies that the protein Fano factor is proportional to the relative noise of the hybrid model. The factor of proportionality is the mean copy number *Ω*/*a* of protein molecules produced per burst. The hybrid model can thus be consistent with a range of different Fano factors depending on the chosen value of the system size. Provided that the value of the system size is fixed to a sufficiently large value, the response of the Fano factor to increasing feedback strength coincides with that of the relative noise (Fig 5, lower panels).

## 10 Summary

We evaluated the noise suppression capabilities of feedback in burst frequency and feedback in decay rate using a hybrid model for bursty protein dynamics. Using a relative noise measure, we systematically related the noise levels of a regulated protein to those of a constitutive protein expressed at the same mean value. It was found that introducing feedback of either kind brings about a decrease in the relative noise. Nevertheless, feedback in decay rate was shown to perform better in suppressing noise, in particular under high-noise and/or strong-feedback conditions.

We identified the relationships between the hybrid model and other modelling frameworks, in particular a deterministic one, based on an ordinary differential equation, and a discrete stochastic framework. The deterministic model is recovered from the hybrid model in the limit of very frequent bursts. The discrete stochastic model reduces to the hybrid model in the limit of large system sizes. Discrete protein distributions estimated by a kinetic Monte Carlo method were found to be in agreement with the continuous distributions provided explicitly by the hybrid framework. The relative noise metric from the hybrid framework was shown to determine the leading order behaviour of the protein Fano factor in the large system size regime.

Overall, our results illustrate the tractability and usefulness of hybrid frameworks in studying non-linear fluctuations in stochastic gene expression.

## Acknowledgement

PB is supported by the Slovak Research and Development Agency under the contract No. APVV-14-0378 and by the VEGA grant 1/0347/18. AS is supported by the National Science Foundation grant ECCS-1711548.

## Appendix A: Small-noise asymptotics

The potential (25), which applies for both regulation types (22) and (27), is a concave function with a single maximum situated at *x* = *x*_0_, where *x*_0_ satisfies the fixed-point equation (24).

For *a* ≫ 1, the most important part of the pdf lies around the maximum *x* = *x*_0_ of the potential. Therefore we use the parabolic approximation in (17) to obtain

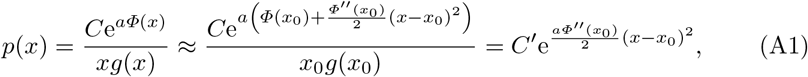

where *C*′ = *C*e^*aΦ*(*x*_0_)^/*x*_0_*g*(*x*_0_) is a constant. The parabolic approximation (A1) implies that at steady state the protein concentration is normally distributed with statistics

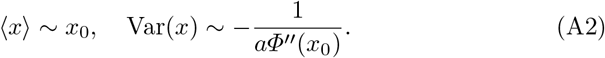

Note that the variance in (A2) is in fact positive since the second derivative of the potential *Φ*(*x*) at the point *x* = *x*_0_ of its maximum is negative.

We see that the (leading-order) approximations (A2) to the protein statistics in the small-noise regime depend on their shared potential (25) but not on the fine differences between the pdfs (26) and (29). Therefore we arrive at a first important conclusion of the present work: feedback in burst frequency and feedback in decay rate are equivalent in terms of control of both mean and noise in the small-noise regime.

Evaluating the second derivative of the potential yields

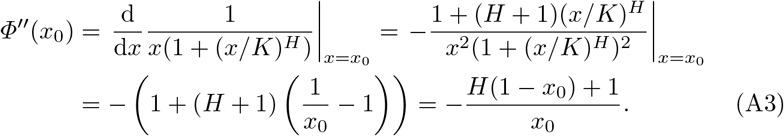

in which we used the fixed-point equation (24) several times. For the relative noise (21) we find

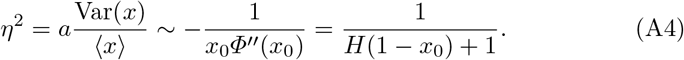

The asymptotic approximation of the relative noise on left-hand side of (A4), which we denote by 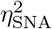, is a decreasing function of *K* which satisfies

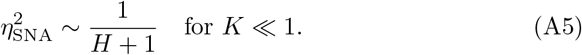

The small-noise asymptotics thus predict that a maximal (*H* + 1)-fold reduction in relative noise can be achieved in the strong-feedback regime using either feedback type.

## 11 Appendix B: Strong-feedback asymptotics

In this Section we develop the relative noise asymptotics for the strong feedback regime *K* ≪ 1. We separately treat feedback in decay rate and refer to literature for treatment of feedback in burst frequency.

### 11.1 Feedback in decay rate

The purpose of this section is to provide asymptotic approximations as *K* ≪ 1 to the integral

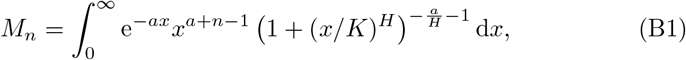

giving the *n*-th moment of the protein pdf (29). In particular, 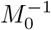 gives the normalisation constant *C* in the protein pdf.

If *K* ≪ 1 and *x* = *O*(1), then *x*/*K* ≫ 1 so that

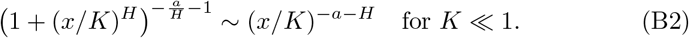

Inserting (B2) into (B1) we find

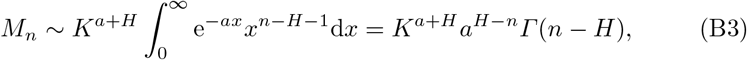

which converges for *n* > *H*. For *n* < *H*, we need to use a different method of approximating the integral (B1).

Substituting (*x*/*K*)^*H*^ = *z* in the integral (B1) yields

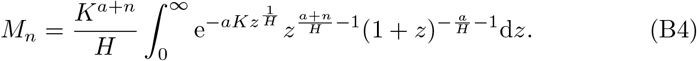

Neglecting the *O*(*K*) term in the exponential in (B4) yields

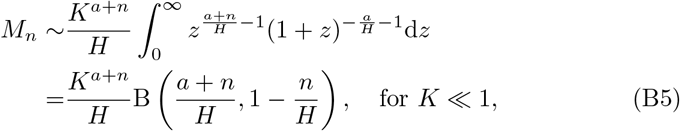

where B(*μ*, *ν*) is the beta function [1]. The right-hand side in (B5) converges for *n* < *H*, which complements the condition for validity of the previous approximation (B5). The nongeneric case *n* = *H* can be treated by method of splitting the integration range [16].

Using the asymptotic approximations (B3) and (B5) in the formula *η*^2^ = *a*(*M*_2_/*M*_1_ − *M*_1_/*M*_0_) for the relative noise, we obtain asymptotic approximations

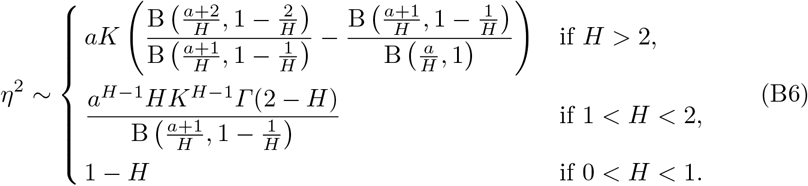

Hence, as 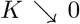, the relative noise decreases to zero linearly if *H* > 2, sub-linearly if 1 < *H* < 2, or tends to a positive constant 1 − *H* if 0 < *H* < 1. High cooperativity in feedback in decay rate thus improves its performance in the strong-feedback regime. Even in the worst case scenario 0 < *H* < 1 in terms of noise control, the limiting value 1 − *H* of relative noise is less than the limiting value 1/(1 + *H*) of the small-noise prediction (A5) for the relative noise.

### 11.2 Feedback in burst frequency

The strong-feedback asymptotic approximation of the relative noise for feedback in burst frequency has been developed in [7] and reads

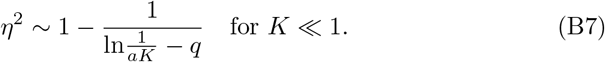

The constant *q* in (B7) is defined by

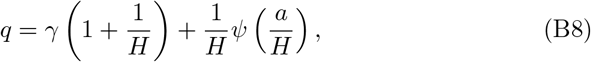

where *γ* is the Euler–Mascheroni constant and *ψ*(*s*) is the digamma function (the logarithmic derivative of the gamma function) [1].

## Appendix C: Skewness of protein distributions

The *n*th central moment is defined by

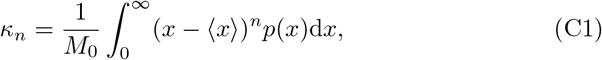

where *p*(*x*) is the protein pdf (26) (feedback in burst frequency) or (29) (feedback in decay rate), *M*_0_ is the normalisation constant (19), and 〈*x*〉 is the protein mean (19).

The skewness of a distribution is defined by

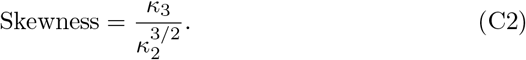

Instead of using absolute skewness values, we report the values of a relative skewness of the steady-state protein distribution (Fig 3 in the Main Text, bottom panels).

The relative skewness compares the skewness in a regulated model to that in a constitutive model with the same mean. The pdf for a referential constitutive model is given by the gamma distribution with shape (mean burst size) 1/*a* and scale *a*〈*x*〉, where 〈*x*〉 is the common value of the mean for the regulated and unregulated models. Since the skewness for the gamma distribution is equal to 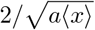, we obtain

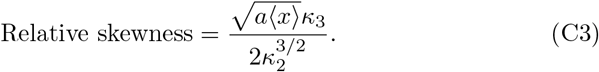

Equation (C3) provides the theoretical basis for the calculations behind the nethermost panels in Fig 3 of the Main Text.

